# Establishment of a transparent soil system to study *Bacillus subtilis* chemical ecology

**DOI:** 10.1101/2022.01.10.475645

**Authors:** Carlos N. Lozano-Andrade, Carla G. Nogueira, Mario Wibowo, Ákos T. Kovács

**Affiliations:** Bacterial Interactions and Evolution Group, DTU Bioengineering, Technical University of Denmark, Kongens Lyngby, Denmark; Natural Product Discovery Group, DTU Bioengineering, Technical University of Denmark, Kongens Lyngby, Denmark

**Keywords:** transparent soil, chemical ecology, *Bacillus subtilis*, fluorescence microscopy, plant-microbe interaction

## Abstract

Bacterial secondary metabolites are structurally diverse molecules that drive microbial interaction by altering growth, cell differentiation, and signaling. *Bacillus subtilis*, a Gram-positive soil-dwelling bacterium, produces a wealth of secondary metabolites, among them, lipopeptides have been vastly studied by their antimicrobial, antitumor, and surfactant activities. However, the natural functions of secondary metabolites in the lifestyles of the producing organism remain less explored under natural conditions, *i.e.* in soil. Here, we describe a hydrogel-based transparent soil system to investigate *B. subtilis* chemical ecology under controllable soil-like conditions. The transparent soil matrix allows the growth of *B. subtilis* and other isolates gnotobiotically and under nutrient-controlled conditions. Additionally, we show that transparent soil allows the detection of lipopeptides production and dynamics by HPLC-MS and MALDI-MS imaging, along with fluorescence imaging of 3-dimensional bacterial assemblages. We anticipate that this affordable and highly controllable system will promote bacterial chemical ecology research and help to elucidate microbial interactions driven by secondary metabolites.

## Introduction

Isolates from the *Bacillus subtilis* species complex have been widely proposed as an alternative to the use of synthetic pesticides in sustainable agriculture. A large set of those soil-dwelling bacteria have been recognized as plant-growth-promoting rhizobacteria (PGPR) since they influence positively plant development via multiple mechanisms (1–3). Production of bioactive secondary (or specialized) metabolites, phytohormones and increased plant nutrient uptake are commonly referred to as putative mode of action involved in plant growth promotion by *Bacilli* (1,4). Several secondary metabolites such as polyketides, terpenes, siderophores, and peptides are thought to be linked to pathogen suppression (biocontrol) or plant defense induction (ISR) (5,6). Production of cyclic lipopeptides (LPs) is the most studied case, although the role of this class has not fully revealed in soil, experiments using mutants impaired in LP secretion suggest the biological significance of LPs in plant growth promotion and biocontrol by *Bacilli* (7–9)

LPs of *B. subtilis* and close relative species are synthesized by non-ribosomal peptide synthase (NRPS), acting as a molecular assembly line that catalyze the amino acid residues incorporation (5,10–12). LPs from *B. subtilis* are grouped into three families: surfactins, iturins and fengycins according to their peptide moieties. They are made up of seven (surfactins and iturins) or ten (fengycins) α-amino acids linked to β-amino (iturins) or β-hydroxyl (surfactin and fengycins) fatty acid (13). LPs encompass a substantial structural diversity associated with a broad spectrum of functionalities (14). To date, role in antagonism toward others organisms, motility, surfaces colonization, and signaling for coordinated growth and differentiation have been described as the main physiological and ecological processes where LPs production is pivotal (15–19). Nevertheless, a mechanistic understanding of how *B. subtilis* secretes LPs, the factor modulating the production and the roles of these compounds under natural conditions remains elusive.

One of the bottlenecks for addressing these questions experimentally is the difficulty to track *in situ* production of LPs and other class of specialized metabolites. Despite the conceptual and analytical chemistry advances detection and quantification of LPs in complex environments such soil and rhizosphere can be particularly difficult, as these amphiphilic compounds can adsorb to soil particles (7). Moreover, occurrence of other compounds can interfere with the detection process as well. A deeper understanding of *B. subtilis* chemical ecology under relevant conditions is crucial for the use of this microbe within sustainable agriculture. Thus, it is a research priority for the *Bacillus* – PGPR community to develop standardized models and methods to elucidate the mechanisms underlying secondary metabolite production *in situ*, to reveal the factors and the experimental variables that define the ecological context of these compounds. To do so, hypothesis-driven research needs to be conducted in highly controlled and adjustable gnotobiotic systems compatible with analytical chemistry and microbiological methods.

Hitherto, phytoagar, peat, mineral substrate (i.e. calcine clay), or hydroponics have been successfully used as gnotobiotic systems to investigate plant-microbe interactions. Many of those systems have been an inexpensive and easy-to-handle alternative for reproducible plant growth under enclosed conditions (20–23). However, none of them has been used for studying secondary metabolites dynamics under soil mimicking conditions. Here, we introduce the hydrogel-based transparent soil developed by Ma and colleagues (24) for plant root phenotyping *in vivo* as gnotobiotic system to investigate *B. subtilis* LPs production and population dynamics under controllable conditions. The transparent soil matrix consists of interconnected pores that are filled with nutrient rich, but diluted cultivation medium, held into spherical beads of hydrogel. We demonstrate that this approach allows the growth of *B. subtilis* and other bacterial strains in axenic, gnotobiotic and nutrient controlled conditions. Lastly, we show that transparent soil allows the detection of LPs production and dynamics by HPLC-MS and MALDI-imaging in addition to fluorescence imaging of 3-dimensional bacterial assemblages.

## Materials and methods

### Bacterial strains used in the study

All the experiments were conducted using the strain *B. subtilis* P5_B1 or its *gfp*-labeled variant P5_B1_*gfp*_ (DTUB38) (9,15). Moreover, the strains *Pedobacter* sp. D749, *Rhodococcus globerulus* D757, *Stenothropomonas indicatrix* D763, and *Chryseobacterium* sp. D764, isolated from the same sampling site of P5_B1, were included in subsequent experiments (25). The strains were routinely maintained in lysogeny broth (LB) medium (LB-Lennox, Carl Roth; 10 g·l^−1^ tryptone, 5 g·l^−1^ yeast extract, and 5 g·l^−1^ NaCl) at 37 °C while shaking at 220 rpm, while microcosm experiments were performed using 0.1× TSB (tryptic soy broth, CASO Broth, Sigma-Aldrich).

### Transparent soil microcosms

All microcosm experiments were carried out using the hydrogel transparent soil previously described by (24). Briefly, a polymeric solution (2.4 g·l^−1^ sodium alginate and 9.6 g·l^−1^ of Phytagel™ (Sigma-Aldrich)) was dropped into stirred solution of 2% CaCl_2_ allowing rapid formation of spherical beads. Subsequently, the beads were soaked into 0.1× TSB as nutrient solution for two hours. After the equilibration period, the excess liquid was drained and 25 g beads were transferred into 50 mL falcon tubes as experimental unit for all the subsequent experiments.

### Bacterial population dynamics on transparent soil microcosm

The population dynamics of *B. subtilis* P5_B1_*gfp*_ and a set of bacterial strains, either growing in pure culture or mixed as bacterial assemblage, was monitored by colony counting, flow cytometry, stereomicroscopy, or culture-independent 16 rRNA profiling on transparent soil microcosms. For P5_B1_*gfp*_, an overnight culture was diluted to an optical density at 600nm (OD_600_) of 0.1 in 0.1× TSB, and 2.5 mL diluted culture were inoculated into transparent soil microcosms. The microcosm was incubated at 21 °C in static conditions. At days 1, 3, 5, 7, and 10, one g of beads was transferred into a 15 mL Falcon tube, diluted in 0.9% NaCl and vortexed for 10 min. The samples were used for cell number estimation via colony-forming unit (CFU) and flow cytometry, and additionally, the growth was inspected by the fluorescence emitted from the beads under stereomicroscopy (Axio Zoom V16 stereomicroscope equipped with a Zeiss CL 9000 LED light source, HE eGFP filter set #38 (excitation at 470/40 nm and emission at 525/50 nm), and an AxioCam MRm monochrome camera (Carl Zeiss, Germany)) at the end sampling point. For colony counting, 100 μL of the sample was serially diluted, spread onto 0.1× TSA (tryptic soy agar, Sigma-Aldrich), and CFU were estimated after 3 days. For quantification of single cells using flow cytometry, the diluted samples were firstly passed through a Miracloth (Millipore) to remove any trace of beads and diluted 1000× in 0.9% NaCl. Subsequently, 1 ml of each sample was transferred to a 2 ml Eppendorf tube and the samples were assayed on a flow cytometer (MACSQuant^®^ VYB, Miltenyi Biotec). Green fluorescent cells were detected using the blue laser (488 nm) and filter B1 (525/50 nm). In addition, non-inoculated beads and 0.1× TSB were used as control to identify background autofluorescence. For each sample, single events were identified from the SSC-H vs. SSC-A plot and gated into GFP vs. SSC-A plot, where GFP positive cells were identified. Similarly, the population dynamics of the strains *Pedobacter* sp. D749, *R. globerulus* D757, *S. indicatrix* D763, and *Chryseobacterium* sp. D764 were monitored on transparent soil microcosm by colony counting. Briefly, overnight cultures of each strain were diluted to an optical density of 0.1 on 0.1× TSB and 2 mL were inoculated into 20 g of transparent soil. At day 1 and 8 post inoculation, the samples were processed following the same procedure described for *B. subtilis* P5_B1_*gfp*_ enumeration.

Lastly, the population dynamic of a bacterial assemblage composed by all five strains was followed using culture-independent bacterial 16S gene profiling. To conduct this, overnight cultures of the five strains were OD adjusted on 0.1× TSB to 1.0 (P5_B1_*gfp*_) and 0.01 (D749, D757, D763, and D764) and mixed on equal volumes. Then, 2.5 mL of the mixture were inoculated into the transparent soil microcosms at incubated at 21°C. At days 1, 3, 5, 8, 11 and 15, the bacterial assemblage genomic DNA was extracted from one g of beads using DNeasy PowerSoil Pro kit (QIAGEN) following the manufacturer’s instructions. The hypervariable regions V3-V4 of the 16S rRNA gene was PCR-amplified using Fw_V3V4 (5’-CCTACGGGNGGCWGCAG-3’) and Rv_V3V4 (5’-GACTACHVGGGTATCTAATCC-3’) primers tagged with eight nucleotides length barcodes reported by Kiesewalter and colleagues (17). The PCR reactions contained 10.6 μL DNase-free water, 12.5 μL TEMPase Hot Start 2x Master Mix, 0.8 μL of each primer (10 μM), and 0.3 μL of 50 ng/μL DNA template. The PCR was performed using the conditions of 95 °C for 15 min, followed by 30 cycles of 95 °C for 30 s, 62 °C for 30 s, 72 °C for 30 s, and finally, 72 °C for 5 min. All V3-V4 amplicons were purified using the NucleoSpin gel and PCR cleanup kit (Macherey-Nagel) and pooled in equimolar ratios. The amplicon pool was submitted to Novogene Europe Company Limited (United Kingdom) for high-throughput sequencing on an Illumina NovaSeq 6000 platform with 2 million reads (2 × 250 bp paired-end reads). The sequence data processing was conducted using the QIIME 2 pipeline (26). The paired-end reads were demultiplexed (cutadapt), denoised and merged using cutadapt (27) and DADA2 (28), respectively. The 16S rRNA reference sequences with a 99% identity criterion obtained from the SILVA database release 132 were trimmed to the V3-V4 region, bound by the primer pair used for amplification, and the product length was limited to 200–500 nucleotides (29). The taxonomy was assigned to the sequences in the feature table generated by DADA2 by using the VSEARCH-based consensus taxonomy classifier (30). Relative species abundance, as population dynamic parameter, was estimated by importing the QIIME 2 artefacts into the R software (4.1) (31) with the package qiime2R and further processed using phyloseq (32) and dplyr (33). All the graphical visualizations were made with ggplot2 (34). At least three replicates were conducted for all experiments.

### Chemicals

All solvents used for UHPLC-HRMS experiments were LC-MS grade (VWR Chemicals); while for metabolites extraction, the solvents were of HPLC grade (VWR Chemicals). Surfactins standard was purchased from Sigma-Aldrich. Surfactin standard stock solutions were prepared in MeOH in concentrations of 0.1, 1, 10, 50, 100, and 500 μg/mL.

### Extraction of secondary metabolites from transparent soil microcosms

For the chemical profiling, 1 g of bead was transferred into 15 mL falcon tube and macerated with 4 mL of isopropylalcohol:EtOAc (1:3, v/v) containing 1% formic acid. Next, the tubes were sonicated for 60 min. The organic solvent was transferred to a new tube, evaporated to dryness under N2, and re-dissolved in 300 μL of methanol for further sonication over 15 min. After centrifugation at 13400 rpm for 3 min, the supernatants were transferred to HPLC vial and subjected to ultrahigh-performance liquid chromatography-high resolution mass spectrometry (UHPLC-HRMS) analysis.

### Secondary metabolite detection by UHPLC-HRMS

UHPLC-HRMS was performed on an Agilent Infinity 1290 UHPLC system with a diode array detector. UV–visible spectra were recorded from 190 to 640 nm. Liquid chromatography of 1 μL extract (or standard solution) was performed using an Agilent Poroshell 120 phenyl-hexyl column (2.1 × 150 mm, 2.7 μm) at 60 °C using of acetonitrile (ACN) and H_2_O, both containing 20 mM formic acid, as mobile phases. Initially, a gradient of 10% ACN/H_2_O to 100% acetonitrile over 15 min was employed, followed by isocratic elution of 100% ACN for 2 min. The gradient was returned to 10% ACN/H_2_O in 0.1 min, and finally isocratic condition of 10% ACN/H_2_O for 2.9 min, at a flow rate of 0.35 mL/min. HRMS spectra were acquired in positive ionization mode on an Agilent 6545 QTOF MS equipped with an Agilent Dual Jet Stream electrospray ion source with a drying gas temperature of 250 °C, drying gas flow of 8 L/min, sheath gas temperature of 300 °C, and sheath gas flow of 12 L/min. Capillary voltage was set to 4000 V and nozzle voltage to 500 V. MS data analysis and processing were performed using Agilent MassHunter Qualitative Analysis B.07.00.

### MALDI Mass Spectrometry Imaging (MSI)

To survey the spatial distribution of LPs via MALDI imaging, 30 mL of agarose 2% were added to each tube and let solidified for 1h at 4 °C. The samples were sectioned at 0.5 cm thickness, and mounted on an IntelliSlides conductive tin oxide glass slide (Bruker Daltonik GmbH) and covered by spraying 1.75 mL of 2,5-dihydrobenzoic acid (DHB) (20 mg/mL in ACN/MeOH/H_2_O (70:25:5, v/v/v)) in a nitrogen atmosphere and dried overnight in a desiccator prior to IMS measurement. The samples were then subjected to timsTOF flex (Bruker Daltonik GmbH) mass spectrometer for MALDI MSI acquisition in positive MS scan mode with 100 μm raster width and a mass range of 100-2000 Da. Calibration was done using red phosphorus. Briefly, a photograph of the sampled on the IntelliSlide was loaded onto Fleximaging software, three teach points were selected to align the background image with the sample slide, measurement regions were defined, and the automatic run mode was then employed. The settings in the timsControl were as follow: Laser: imaging 100 μm, Power Boost 3.0%, scan range 26 μm in the XY interval, and laser power 90%; Tune: Funnel 1 RF 300 Vpp, Funnel 2 RF 300 Vpp, Multipole RF 300 Vpp, isCID 0 eV, Deflection Delta 70 V, MALDI plate offset 100 V, quadrupole ion energy 5 eV, quadrupole loss mass 100 m/z, collision energy 10 eV, focus pre TOF transfer time 75 μs, pre-pulse storage 8 μs. After data acquisition, the data were analyzed using SCiLS software.

### Root colonization assay

The transparent soil microcosm was assayed for supporting plant growth and root colonization by *B. subtilis* in the early stages of tomato seedlings development (*Solanum lycopersicum* L., Maja Buschtomato, Buzzy Seeds, NL). Seeds were surface sterilized in Eppendorf tubes by shaking in an orbital mixer for 10 min in 1.5 mL of 2% sodium hypochlorite. Afterward, seeds were washed five times in sterile MiliQ water alternating centrifugation and removal of liquid solution. Then, 10 seeds were germinated on 15% agar for 3 days. Subsequently, seedlings were soaked on a P5_B1_*gfp*_ bacterial solution (OD_600_ = 0.1) for 10 minutes and placed in LEGO brick boxes containing 50 g of beads (35). The root colonization was tracked by confocal laser scanning microscopy imaging (CLSM) as described previously (15,36–39). Colonized roots were washed twice with sterile ddH_2_O and placed onto microscope slides. Images were captured in a Leica TCS SP8 microscopy. Green fluorescent reporter excitation was performed at 488 nm, while the emitted fluorescence was recorded at 520/23 nm. For generating multilayer images, Z-stack series with 1μm steps were acquired and processed with the software Fiji (40).

## Results

### A transparent soil microcosm for studying *B. subtilis* and other bacterial species chemical ecology

We established a transparent soil microcosm for studying microbial interactions under axenic and soil mimicking conditions motivated by the alginate bead-based system described by Ma and colleagues (24). The developed system can be used to measure both population dynamic parameters and metabolites production as a proxy for bacterial establishment in the system. We aimed to determine growth and viability using plate colony counting, flow cytometry, microscopy, and 16S rRNA gene profiling on pure cultures or when the strains are co-cultivated in the microcosm. Additionally, LPs produced by *B. subtilis* was detected and quantified using UHPLC-HRMS and MSI that aims to facilitate the study of bacterial interactions driven by this class of compounds in a controllable system (Fig. 1A). Therefore, the depicted system allows research on microbial community chemical ecology under defined laboratory conditions.

**Fig. 1.**
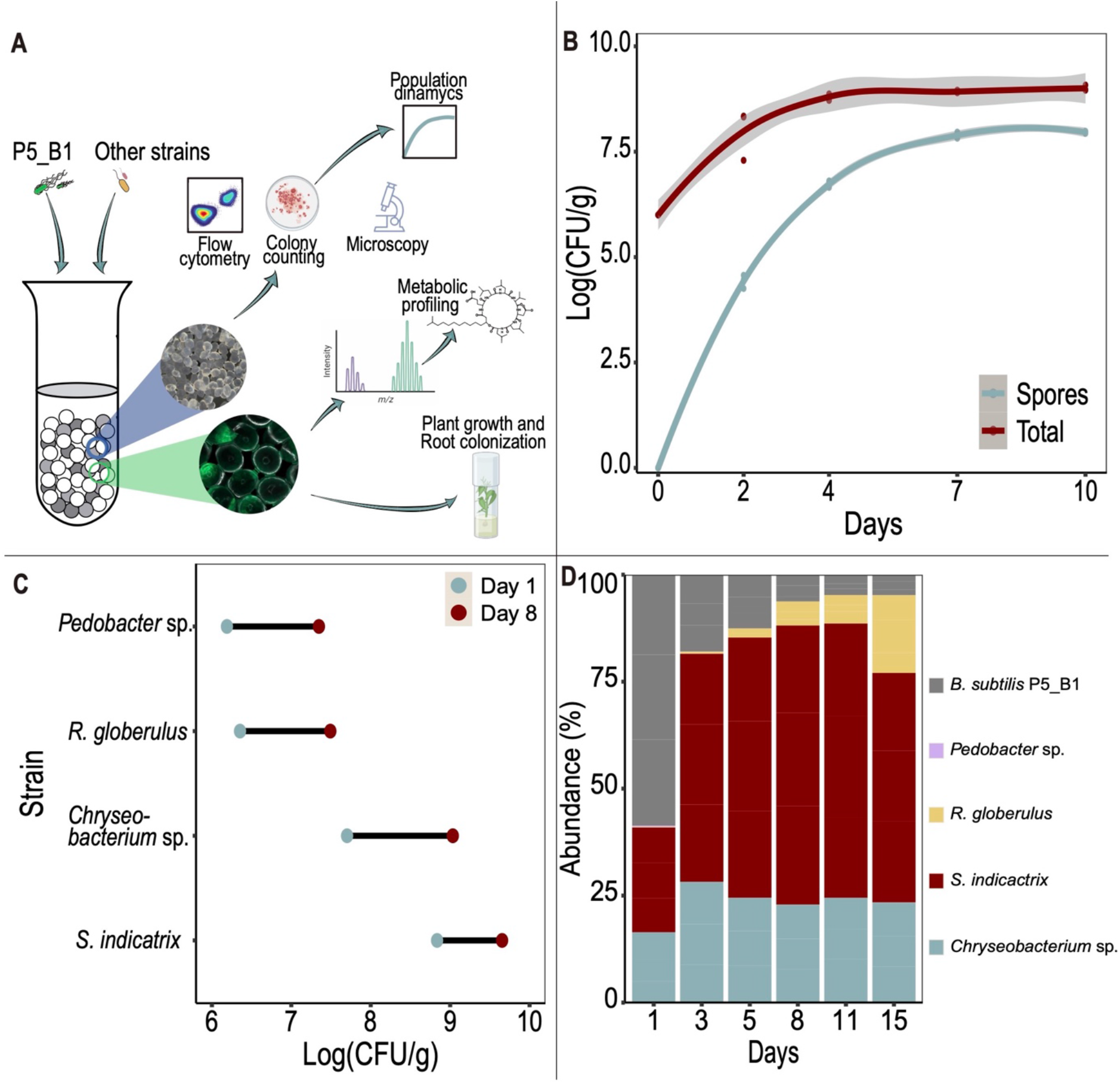
A transparent soil microcosm supports the growth of *B. subtilis* and other bacterial strains. (A) Overview of the experimental approach followed with the transparent soil microcosm. (B) Changes in *B. subtilis* P5_B1 populations (spore and total number of cells) on transparent soil microcosm were monitored as CFU/g over time (*n* = 3). The solid lines represent adjusted curve from a generalized model using the function *stat_smooth* in R. The gray area represents the dispersion given as confidence interval at 95%. (C) Endpoint population changes of four bacterial species on the soil microcosms. Population growth after day 1 and day 8 post inoculation were estimated by CFU/g (*n* = 3). (D) Culture-independent population dynamic estimation. A taxonomic summary showing the relative abundance of the five bacterial species inoculated into the transparent soil over 15 days (*n* = 3).

### A transparent soil microcosm supports the growth of *B. subtilis* and other bacterial isolates

To demonstrate the capability of the transparent soil to support bacterial growth and development, we followed the *on-microcosm* population dynamics of P5_B1 labeled with a constitutively expressed green fluorescent protein in three replicates using plate count and flow cytometry during 10 days (Fig. 1). Both the total number of cells and spores increased exponentially for roughly 4 days followed by a plateau until the last sampling point at day 10 post-inoculation reaching around 10^9^ CFU g^−1^ maximum carrying capacity. Sporulation is a key phenotypic developmental trait to consider in *Bacillus* chemical ecology, given the impact of dormant spore may have on fitness and secondary metabolites overall production. In our single species microcosm experiments, the spore population varied from around 55%, at day two, to a maximum level of roughly 88% after 10 days (Fig. 1B).

Similarly, as revealed by plate colony count, the transparent soil microcosms can sustain the growth of other soil-derived bacterial isolates. Here, determination of growth properties confirmed that all the strains grew and increased their population in at least two orders of magnitude compared to the initial inoculation ratio (~10^6^ CFU g^−1^) (Fig 1 C).

Additionally, we surveyed whether the microcosms is suitable to grow a bacterial assemblage and estimate its composition using a culture-independent method that relies on environmental DNA extractions. As we expected, given the low complexity of the transparent soil compared to soil, we obtained a high quality and quantity of environmental DNA from the system (>220 ng/μL) allowing us to determine the bacterial composition over 15 days by 16 rRNA gene sequencing. At day 1, we detected all the five strains at different proportions, being P5_B1*gfp* the most abundant strain because of its initial inoculation ratio (1:100). Over time, the bacterial assemblage experienced strong composition changes, whereas *S. indicatrix* and *R. globerulus* substantially increased their population, P5_B1*gfp* was downsized to around 10% of its abundance, and *Pedobacter* sp. was totally outcompeted; at the end of the experiment, *S. indicatrix* and *Chryseobacterium* sp. were the most abundant taxa of the assemblage (Fig. 1D).

### Surfactin and plipastatin are produced at detectable levels in the transparent soil microcosms

One bottleneck in chemical ecology research on *Bacilli* is imposed by the difficulties to detect and quantify LPs and other secondary metabolites in their niche where these are naturally produced, limiting our understanding of how those compounds impact the ecology of producers and interacting species. Therefore, to shed light on the qualitative productions of LPs in our experimental system, we monitored the metabolic profile of P5_B1 using UHPLC-HRMS, targeting compounds with *m/z* values between 1.000 and 1.600, which was the typical *m/z* range for *Bacillus* LPs detection (41). In the axenic cultures of P5_B1, we detected the multiples isoforms belonging to the surfactin and plipastatin families, being surfactin C15 and plipastatin B C17, as the main components of LPs mixture (Fig. 2, Table 1). To corroborate the HPLC-MS findings and dissect the spatial distribution of surfactin and plipastatin in our experimental system, we tracked the LPs production by MALDI imaging mass spectrometry. The isoforms surfactin C15 and plipastatin B C17 were massively detected in all bead section surveyed, confirming that those compounds are diffusible in the matrix (Fig. 3).

**Fig. 2.**
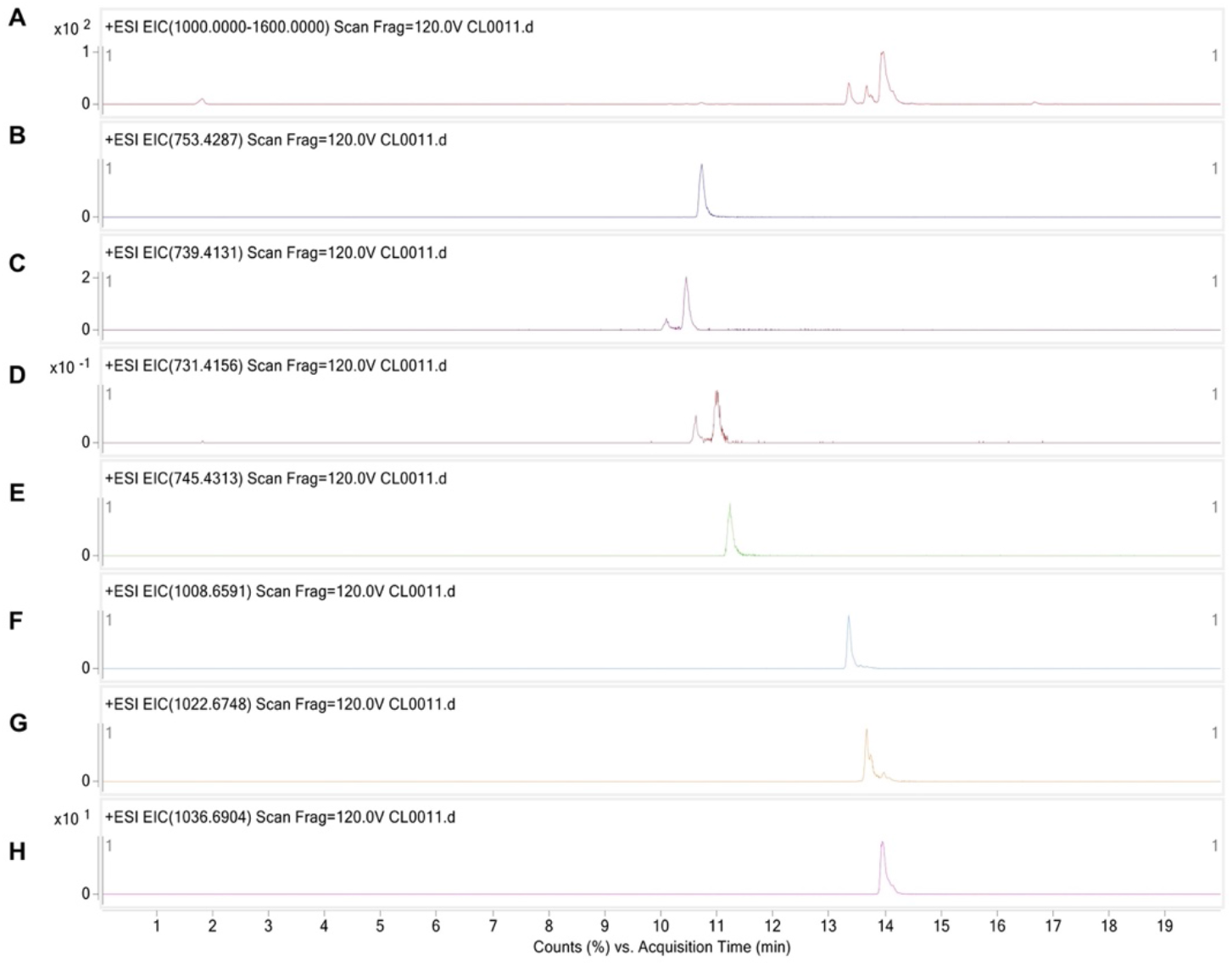
Extracted ion chromatograms (EIC) for **A)** lipopeptides detection (*m/z* 1000-1600), **B)** Plipastatin B C_17_ (*m/z* 753.4287 ± 10 ppm), **C)** Plipastatin B C_15_/Plipastatin A C_17_ (*m/z* 739.4131 ± 10 ppm), **D)** Plipastatin B C_14_/Plipastatin A C_16_ (*m/z* 731.4156 ± 10 ppm), **E)** Plipastatin B C_16_ (*m/z* 745.4313 ± 10 ppm), **F)** Surfactin-C_13_ (*m/z* 1008.6591 ± 10 ppm), **G)** Surfactin-C_14_ (*m/z* 1022.6748 ± 10 ppm), **H)** Surfactin-C_15_ (*m/z* 1036.6904 ± 10 ppm).

**Table 1.** Identified lipopeptides.

| Retention time (min) | *m/z* | ID | Formula |
| --- | --- | --- | --- |
| 10.73 | 753.4292 [M+2H]<sup>2+</sup>,<br>1505.8507 [M+H]<sup>+</sup> | Plipastatin B C<sub>17</sub> | C<sub>75</sub>H<sub>116</sub>N<sub>12</sub>O<sub>20</sub> |
| 10.08 | 739.4141 [M+2H]<sup>2+</sup> | Plipastatin B C<sub>15</sub> | C<sub>73</sub>H<sub>112</sub>N<sub>12</sub>O<sub>20</sub> |
| 10.45 | 739.4127 [M+2H]<sup>2+</sup> | Plipastatin A C<sub>17</sub> | C<sub>73</sub>H<sub>112</sub>N<sub>12</sub>O<sub>20</sub> |
| 10.62 | 731.4169 [M+2H]<sup>2+</sup> | Plipastatin B C<sub>14</sub> | C<sub>73</sub>H<sub>112</sub>N<sub>12</sub>O<sub>19</sub> |
| 11.02 | 731.4148 [M+2H]<sup>2+</sup> | Plipastatin A C<sub>16</sub> | C<sub>73</sub>H<sub>112</sub>N<sub>12</sub>O<sub>19</sub> |
| 11.24 | 745.4305 [M+2H]<sup>2+</sup>,<br>1489.8552 [M+H]<sup>+</sup> | Plipastatin B C<sub>16</sub> | C<sub>75</sub>H<sub>116</sub>N<sub>12</sub>O<sub>19</sub> |
| 13.35 | 1008.6593 [M+H]<sup>+</sup>,<br>1030.6409 [M+Na]<sup>+</sup> | Surfactin-C<sub>13</sub><sup>a</sup> | C<sub>51</sub>H<sub>89</sub>N<sub>7</sub>O<sub>13</sub> |
| 13.66 | 1022.6764 [M+H]<sup>+</sup>,<br>1044.6580 [M+Na]<sup>+</sup> | Surfactin-C<sub>14</sub><sup>a</sup> | C<sub>52</sub>H<sub>91</sub>N<sub>7</sub>O<sub>13</sub> |
| 13.97 | 1036.6910 [M+H]<sup>+</sup>,<br>1058.6728 [M+Na]<sup>+</sup> | Surfactin-C<sub>15</sub><sup>a</sup> | C<sub>53</sub>H<sub>93</sub>N<sub>7</sub>O<sub>13</sub> |

<sup>a</sup> Validated by LCMS of authentic standard

**Fig. 3.**
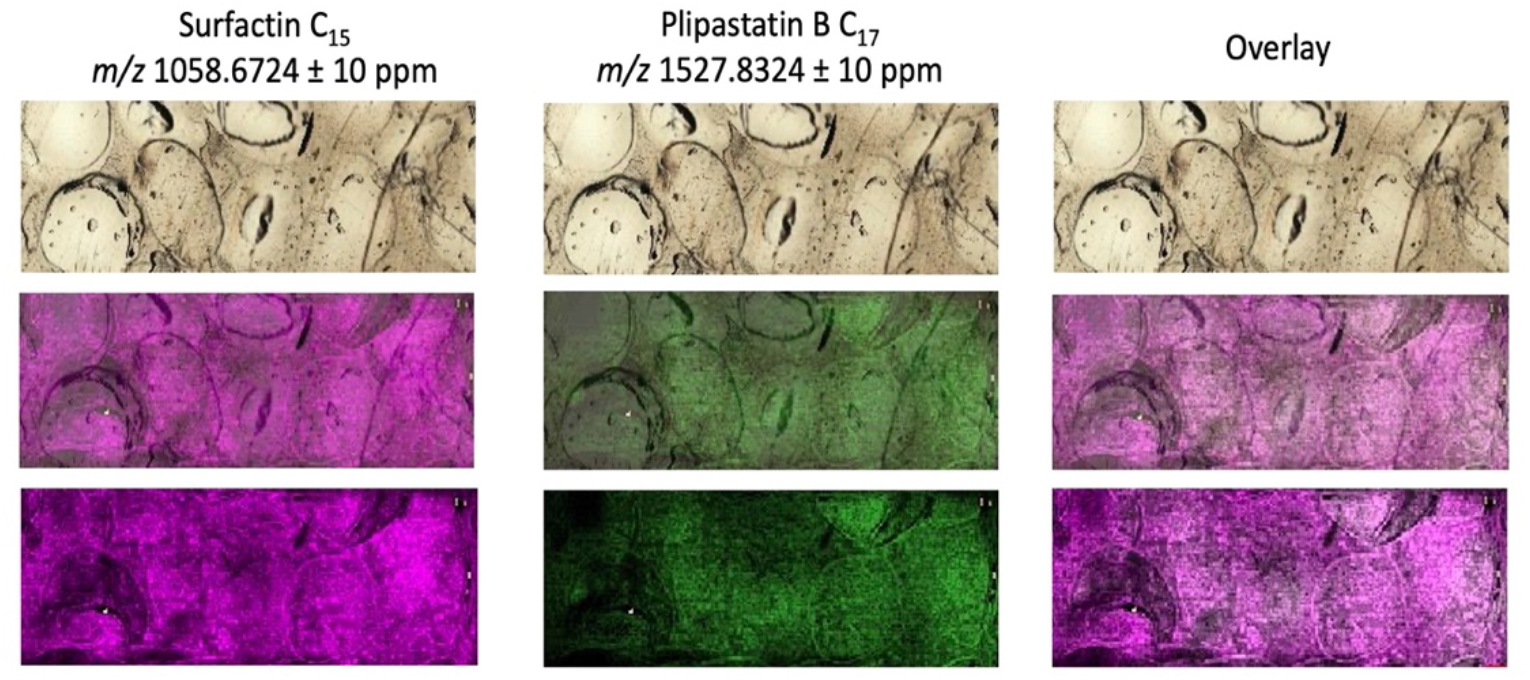
Positive ion mode MALDI imaging analysis of *B. subtilis* cultured in the hydrogel matrix. For the MALDI-MSI, the hydrogel matrix was covered with 2% agarose and incubated for 2h at 4°C. Then, the plugs were cross-sectioned before the MALDI-MSI inspection. The ion intensity is reflected by the intensity of colors.

### The transparent soil microcosm allows plant growth and serves as gnotobiotic system for studying *B. subtilis* root colonization

To examine whether the microcosm can support plant growth and serve as model for *B. subtilis* root colonization assays on gnotobiotic conditions, pre-germinated tomato seeds were inoculated with a bacterial suspension and placed on the plant cultivation box based on LEGO assemblies (35). Overall, the plants emerged after 4 days post-inoculation and, during two weeks of growth, the leaves appeared healthy and dark green. Noticeably, the roots system grew profusely allowing subsequent inspection of P5_B1*gfp* colonization under CLSM. Here, P5_B1*gfp* formed robust biofilm on the root and fluorescent cells were detected up to 10 days post inoculation, suggesting that the system can be used for interrogating the role of LPs in root colonization in early stages (Fig. 4).

**Fig. 4.**
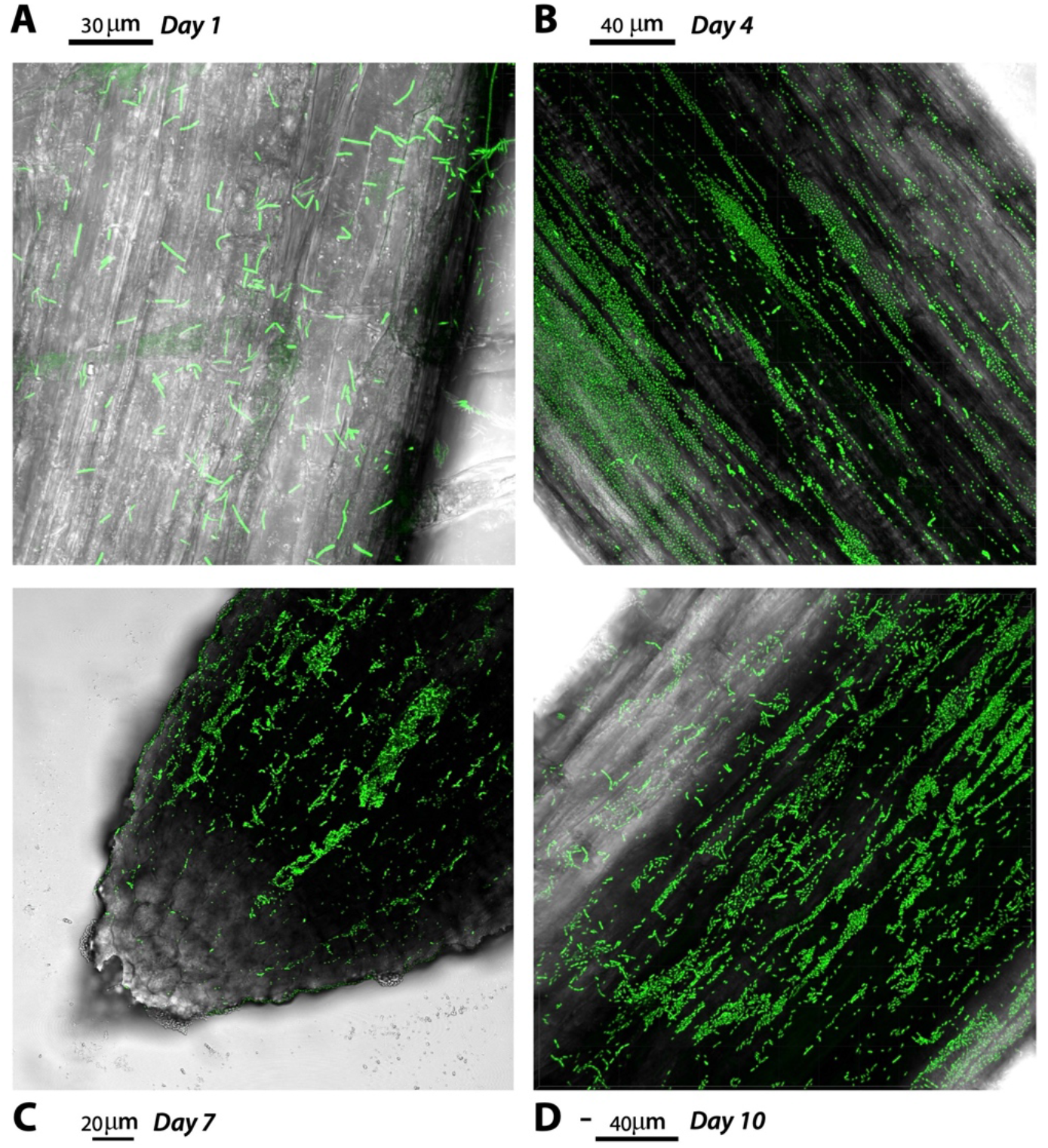
*B. subtilis* P5_B1*gfp* root colonization patterns on tomato seedlings.

## DISCUSSION

Secondary metabolite production is intended as one of the main drivers of microbial interaction between *Bacilli* and other microorganisms (42,43). Their relevance on such processes has been elegantly demonstrated for a large set of secondary metabolites in different bacterial strains, revealing roles on cell differentiation, antagonisms and signaling (8). However, studies shedding light on the role of those SM under natural conditions or in model systems mimicking natural niches remain scarce.

In this proof-of-concept study, we demonstrated that a hydrogel matrix developed by Mao and colleagues can be used as inexpensive and highly controllable microcosms for studying *B. subtilis* chemical ecology combining microbiological and chemical methods. The system, despite its simplicity, arises as an alternative to overcome the technical bottlenecks imposed by soil complexity in terms microbial diversity and metabolites quantification.

As we demonstrated here, the transparent soil microcosms can be used for addressing a variety of questions in microbial ecology under simplified conditions otherwise hard to test in natural environments (*i.e.* soil). First, the system can be exploited to describe bacterial population dynamics either under axenic cultures (monoculture) or growing as a synthetic community using classic colony counting, flow cytometry, and 16s rRNA profiling. In addition, the system’s capability to support other microorganism’s growth allows dissecting fundamental questions about the consequences of community diversity on *Bacilli* success and SM production. For instance, revealing the importance of richness (number of interacting species) and structure (how the members contribute to the overall community performance) on bacterial interaction where SM production may be relevant.

Tracking LPs production in complex environments such as soil and rhizosphere has been extremely difficult and represent one of the main limitations to interrogate the biological activities of LPs under natural conditions. Here, we demonstrated that the described system allows exploring secondary metabolites dynamics looking at both spatial and temporal distribution by coupling HPLC-MS and MALDI imaging, overcoming limitations associated with extraction, detection, and quantification of LPs in soil. Likewise, untargeted metabolic analysis could be performed in order to detect unknown compounds occurring after *B. subtilis* or other microorganisms growing in the matrix, since interferences caused by the large amount of organic matter are not present in the system (8,44,45).

As Mao and colleagues demonstrated, the hydrogel transparent soil produces field-relevant root phenotypes in *Glycine max* (24). Therefore, we interrogated whether bacterial-inoculated tomato seedlings could be grown on the described system and also studied root colonization pattern by *B. subtilis*. As we expected, the microcosms provided optimal controlled growth conditions for tomato and the plant-associated bacterium *B. subtilis* facilitating the study of host-microbe interactions possibly influenced by secondary metabolites in the rhizosphere.

Overall, the system offers several experimental advantages and a high degree of customization. Nevertheless, as is true for lab-scale simplified experiments, there are several limitations compared to assays conducted in natural soil. First, the absence of organic matters and/or soil-like structure might alter microbial interactions and community assembly patterns (46). Moreover, not all known secondary metabolites potentially produced by *B. subtilis* were detected using HPLC-MS and MALDI imaging. To date, it has been a common limitation on bacterial chemical ecology studies *in situ*, that will be overcome as analytical chemistry and soil-mimicking systems advance.

In conclusion, we have established a novel transparent soil microcosm that enables us to examine microbial interactions where secondary metabolites and especially, lipopeptides could be pivotal. It is a simplified soil-mimicking matrix that allows registering the growth of microbes, plants and detecting diverse metabolites under controllable conditions. The system, despite being a reductionist approach but still retaining some degree of soil-like properties, can be helpful to disentangle the inherent complexity of interactions that occur in natural soil.

## ACKNOWLEDGMENTS

This project was supported by the Danish National Research Foundation (DNRF137) for the Center for Microbial Secondary Metabolites. Funding from Novo Nordisk Foundation for the infrastructure “Imaging microbial language in biocontrol (IMLiB)” (NNFOC0055625) and via the project INTERACT (grant number NNF19SA0059360) is acknowledged.

## AUTHOR CONTRIBUTIONS

CNLA and ATK conceived and designed the study. CNLA, CGN and MW conducted the experiments, and CNLA and MW analyzed data. CNLA and ATK wrote the manuscript; all authors approved the manuscript.

